# Targeted CRM1-inhibition perturbs leukemogenic NUP214 fusion proteins and exerts anti-cancer effects in leukemia cell lines with *NUP214* rearrangements

**DOI:** 10.1101/2020.04.23.057257

**Authors:** Adélia Mendes, Ramona Jühlen, Valérie Martinelli, Birthe Fahrenkrog

**Author notes:** corresponding author: Birthe Fahrenkrog.

## Abstract

Chromosomal translocations fusing the locus of nucleoporin *NUP214* each with the proto-oncogenes *SET* and *DEK* are recurrent in, largely intractable, acute leukemias. The molecular basis underlying the pathogenesis of SET-NUP214 and DEK-NUP214 are still poorly understood, but both chimeras inhibit protein nuclear export mediated by the ß-karyopherin CRM1. In this report, we show that SET-NUP214 and DEK-NUP214 both disturb the localization of proteins essential for nucleocytoplasmic transport, in particular for CRM1-mediated protein export. Endogenous and exogenous SET-NUP214 and DEK-NUP214 form nuclear bodies. These nuclear bodies disperse upon targeted inhibition of CRM1 and the two fusion proteins re-localize throughout the nucleoplasm. Moreover, SET-NUP214 and DEK-NUP214 nuclear bodies reestablish shortly after removal of CRM1 inhibitors. Likewise, cell viability, metabolism, and proliferation of leukemia cell lines harboring SET-NUP214 and DEK-NUP214 are compromised by CRM1 inhibition, which is even sustained after clearance from CRM1 antagonists. Our results indicate CRM1 as a possible therapeutic target in NUP214-related leukemia. This is especially important, since no specific or targeted treatment options for NUP214 driven leukemia are available yet.

## Introduction

Chromosomal translocations involving the nucleoporin *NUP214* have been described in *de novo* and therapy-related acute myeloid leukemia (AML) as well as acute lymphoblastic leukemia (ALL). NUP214-related malignancies are frequently associated with poor treatment response and poor prognosis^1-6^. The fusion proteins SET-NUP214 [del(9)(q34.11q34.13)] and DEK-NUP214 [t(6;9)(p23;q34)] result from the fusion of the almost entire SET and DEK proteins with the C-terminal part of NUP214^1, 7, 8^. NUP214 is an integral part of the nuclear pore complex (NPC) and it plays important roles in nuclear export mediated by chromosomal region maintenance 1 (CRM1, or exportin 1/XPO1)^9, 10^. CRM1 is the major nuclear export receptor for proteins and ribonucleoprotein (RNP) complexes carrying a characteristic nuclear export signal (NES)^11, 12^. NUP214 functions as a terminal docking site for CRM1 nuclear export complexes on the cytoplasmic side of NPCs and depletion of NUP214 results in nuclear accumulation of NES-containing cargoes^13-16^.

The C-terminal phenylalanine-glycine (FG) repeat domain of NUP214 exhibits multiple CRM1-binding sites, which are preserved in SET-NUP214 and DEK-NUP214^16-19^. In fact, both fusion proteins can bind CRM1 and its co-factor, the small GTPase Ran, and inhibit the nuclear export of NES-containing proteins and RNPs^17, 18^. Targeted CRM1 inhibition by small molecule antagonists has become an appealing anti-cancer strategy, for both solid and hematologic malignancies^20-24^. Leptomycin B (LMB), a fungal metabolite from *Streptomyces* spp, was the first identified small molecule inhibitor specifically targeting CRM1^25^. LMB has potent anti-cancer activity, but its application in patients was withdrawn after a single phase I clinical trial because of its low efficiency and high toxicity^26^. Selective inhibitors of nuclear export (SINEs) comprise a novel class of CRM1 antagonists with anti-cancer properties both *in vitro* and *in vivo*^20-23, 27-29^. Indeed, the SINE compound KPT-330 is currently tested in phase 2/3 clinical trials for a wide variety of cancers, including leukemia and other hematologic malignancies^30^. The anti-cancer effects of CRM1 inhibitors are based on the induction of cell death by apoptosis and on cell cycle arrest due to activation of the transcriptional programs of tumor suppressor genes, such as *TP53, RB1* and *FOXO*-related tumor suppressors^20, 21, 31^. Despite the functional proximity between NUP214 and CRM1-mediated nuclear export and the functional relevance of CRM1 in some leukemia, the impact of CRM1 inhibition in the context of NUP214-related leukemia has not been studied yet.

Here we used patient-derived leukemia cell lines expressing endogenous SET-NUP214 and DEK-NUP214 to address the anti-cancer potential of CRM1 inhibition in NUP214-rearranged leukemia. We report that CRM1 inhibition by LMB or the SINE compound KPT-185 is sufficient to disturb the localization of endogenous SET-NUP214 and DEK-NUP214, coinciding with reduced cell viability and proliferation. CRM1 inhibition reduces cell viability and metabolic activity in a sustained manner after drug withdrawal. However, after drug removal, proliferation and SET-NUP214 and DEK-NUP214 nuclear bodies are restored. Our data suggest that CRM1 constraint is an interesting candidate for the development of an anti-cancer therapeutic approach in NUP214-related leukemia, for which no efficient targeted therapy has been developed so far.

## Material and Methods

### Cell lines

LOUCY (T-cell leukemia), MEGAL (acute megakaryoblastic leukemia), FKH-1 (acute myelocytic leukemia), OCI-AML1 (acute myeloid leukemia), and MOLM-13 (acute myeloid leukemia) cell lines were purchased from the Leibniz Institute - DSMZ-German Collection of Microorganisms and Cell Cultures GmbH (Braunschweig, Germany). HCT-116 cells were a gift from Dr. Denis Lafontaine (Institute of Molecular Biology and Medicine, Université Libre de Bruxelles, Gosselies, Belgium). Cell lines were maintained as detailed in the *Online Supplementary Methods.*

### Plasmids and Transfections

To generate SET-NUP214-GFP, total RNA was extracted from LOUCY cells using the High Pure RNA Isolation Kit (Roche Life Sciences, Basel, Switzerland) according to the manufacturer’s instructions. cDNA synthesis was performed by reverse transcription-PCR. Cloning of SET-NUP214 is described in the *Online Supplementary Methods*.

The pENTR1-DEK-NUP214 plasmid was a gift from Dr. Martin Ruthardt (Cardiff University, UK) and was subcloned into the peZY-EGFP destination vector using the Gateway™ LR clonase™ enzyme mix (Invitrogen, Merelbeke, Belgium), as described in the *Online Supplementary Methods.*

### Immunofluorescence of suspension cells

Leukemia cells were seeded at 0.8×10^6^ cells/ml and grown for 24 h - 48 h. Cells were fixed and processed for immunofluorescence as detailed in the *Online Supplementary Methods.*

### Immunofluorescence of adherent cells

HCT-116 cells were seeded on polylysine-coated glass coverslips 24 h prior to transfection. After transfection, cells were fixed in 2% formaldehyde and processed for immunofluorescence as detailed in the *Online Supplementary Methods.*

### 1,6-hexanediol treatment

LOUCY cells were seeded at 0.8×10^6^ cells/ml 24 h prior to treatment. Cells were then incubated with 5% 1,6-hexanediol (Sigma-Aldrich, Overijse, Belgium) for different time-points. After treatment, cells were washed and processed for immunofluorescence as detailed in the *Online Supplementary Methods*.

### Inhibition of CRM1-mediated nuclear export

Cells were seeded at 0.8×10^6^ cells/ml for 24 h. Next, cells were treated with 1 µM KPT-185 (Selleck Chemicals, Munich, Germany) or 20 nM LMB (Enzo Life Sciences, Brussels, Belgium). Cell viability and proliferation was assessed by (i) Trypan Blue exclusion dye assay, (ii) the cell proliferation reagent WST-1 (Roche Life Sciences) according to the manufacturer’s instructions, and (iii) flow cytometry using FITC-Ki-67 (BD Biosciences, CA, USA). The detailed protocols are described in the *Online Supplementary Methods.*

### Statistics

Experiments were performed at least three times and the results represent the mean ± SEM for three independent biological replicates. Plots were generated and statistical analysis was performed using GraphPad Prism (Version 5.01; GraphPad Software Inc., CA, USA). Statistical differences were calculated by one-way analysis of variance (ANOVA). During evaluation of the results a confidence interval α of 95% and p values lower than 0.05 were considered as statistically significant. Significance levels are represented as *(p<0.05), **(p<0.01) or **(p<0.001).

## Results

### Leukemogenic NUP214 fusion proteins locate to nuclear bodies in patient-derived cells

We first determined the localization of NUP214 fusion proteins in different patient-derived leukemia cell lines with anti-NUP214 antibodies and immunofluorescence microscopy. LOUCY and MEGAL cells express SET-NUP214 and in both cell lines SET-NUP214 located to the nuclear rim and to nuclear bodies (Figure 1A), consistent with previous results^17, 32^. FKH-1 cells harbor DEK-NUP214, which localized to smaller nuclear bodies as compared to SET-NUP214 (Figure 1A)^33^. Similar localizations for GFP-tagged versions of SET-NUP214 and DEK-NUP214 were observed in transiently transfected HCT-116 cells (Figure 1B). In FKH-1 cells, NUP214 antibodies were also detected at the nuclear rim, which likely corresponds to endogenous NUP214 rather than to the fusion protein, as DEK-NUP214-GFP in HCT-116 was not detected at NPCs (Figure 1 A-B). In OCI-AML1 and MOLM-13 cells, which do not express NUP214 fusion proteins, NUP214 staining displayed the typical punctate pattern of nucleoporins at the nuclear rim (Figure 1A).

**Figure 1:**
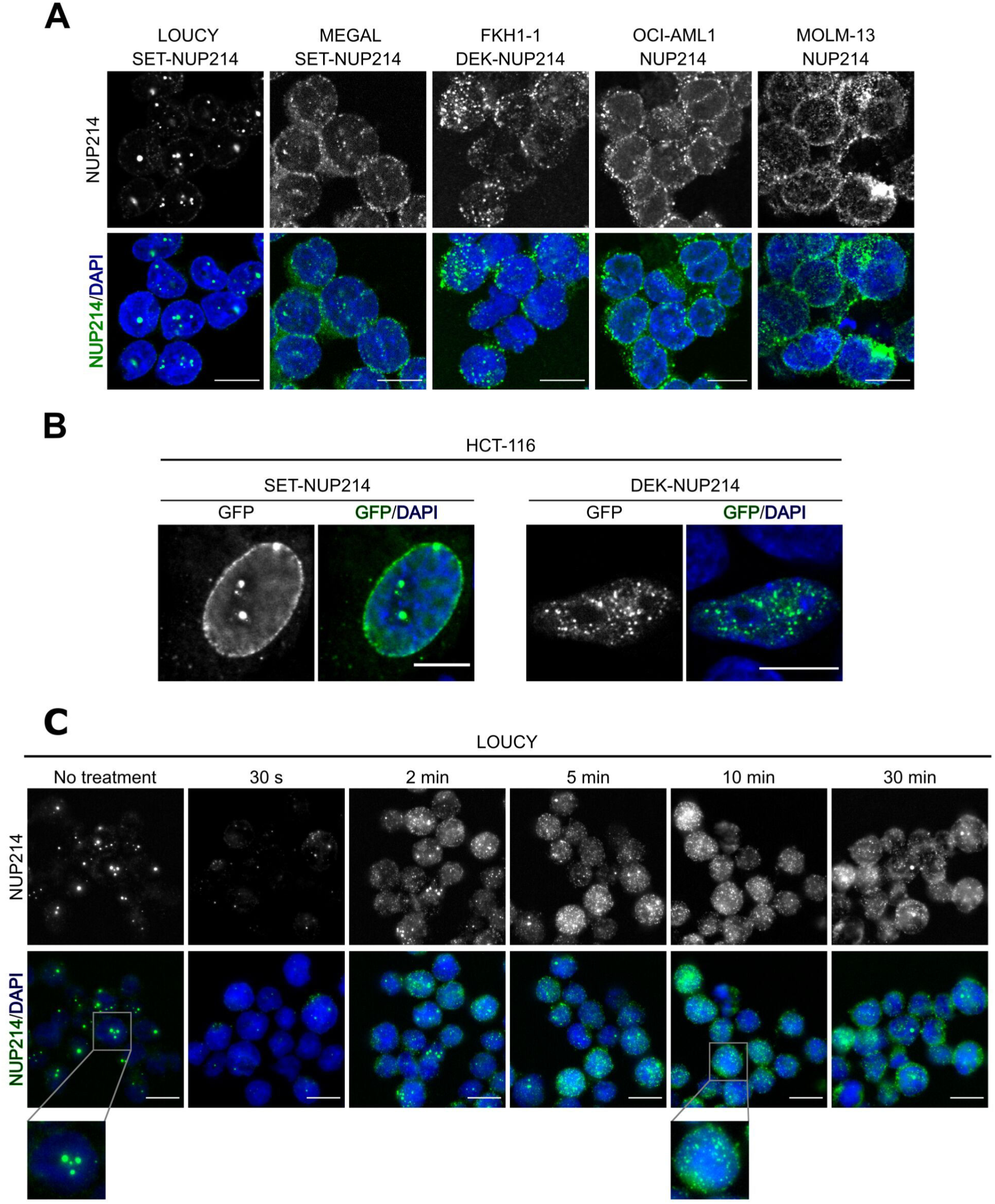
NUP214 fusion proteins localize to distinct nuclear bodies **(A)** Cellular distribution of NUP214 in distinct leukemia cell lines. Foci correspond to nuclear bodies formed by SET-NUP214 (LOUCY and MEGAL) or DEK-NUP214 (FKH-1). **(B)** HCT-116 cells transfected with SET-NUP214-GFP or DEK-NUP214-GFP. **(C)** LOUCY cells were treated with 5% 1,6-hexanediol for the indicated time points. The presence of nuclear bodies was evaluated by immunofluorescence of NUP214 (green). DNA was visualized with DAPI (blue). Shown are representative epifluorescence (A, C) and confocal images (B). Scale bars, 10 µm.

The FG domains of nucleoporins are, due to their amino acid composition, intrinsically disordered and exhibit variable degrees of cohesiveness, which is important for the maintenance of the NPC permeability barrier^34-36^. 1,6-hexanediol (HD) is a mild alcohol that interferes with hydrophobic interactions established between FG repeats^34^, thereby disrupting the NPC permeability barrier ^37, 38^. To address the potential role of FG repeat cohesion in the formation of NUP214 nuclear bodies, we treated LOUCY cells, which have the most prominent nuclear bodies, with 5% HD and monitored SET-NUP214 nuclear bodies over time by immunofluorescence microscopy. As shown in Figure 1C, untreated cells represented with prominent SET-NUP214 nuclear bodies of different size and frequency. These SET-NUP214 nuclear bodies started to dissolve as early as 2 min after HD treatment and vanished after 10 min HD exposure, as indicated by a largely spread and homogeneous distribution of SET-NUP214 throughout the nucleus (Figure 1C). Hence, we concluded that the formation of SET-NUP214 nuclear bodies depends, at least in part, on cohesive interactions between the FG repeats of NUP214.

### NUP214 fusions accumulate the nuclear export factor CRM1 in their nuclei and perturb the localization of endogenous nucleoporins

We have previously shown that SET-NUP214 in transfected cells interacts with the nuclear export factor CRM1 and that this interaction is enhanced in the presence of RanGTP^17^. Accordingly, we found that SET-NUP214 nuclear bodies accumulate CRM1 in LOUCY (Figure 2A) as well as endogenous CRM1 and exogenous Ran-RFP in transfected HCT-116 cells (Figure 2 B-C). We made similar observations for endogenous and exogenous DEK-NUP214, however, due to the smaller size of the nuclear foci, the co-localization of the fusion protein with Ran-RFP was less obvious than observed for SET-NUP214 (Figure 2 A-C). In contrast, CRM1 was enriched at the nuclear membrane in OCI-AML1 cells (Figure 2A) and localized throughout the nucleus and cytoplasm in HCT-116 cells expressing GFP alone (Figure 2B), reflecting the normal range of CRM1 distribution in cells (Human Protein Atlas, available from http://www.proteinatlas.org)^39^. To reinforce the notion that NUP214 fusion proteins preferentially bind CRM1-RanGTP nuclear export complexes, we co-expressed SET-NUP214-GFP and DEK-NUP214-GFP in HCT116 cells with RFP-tagged versions of Ran^17^. We employed RanQ69L (RanQ69L-RFP), a non-hydrolyzable mutant of Ran, and RanT24N (RanT24N-RFP), which is resistant to GTP loading and nuclear export complex formation ^40^. As expected, both NUP214 fusion proteins sequestered RanQ69L-RFP, but not RanT24N-RFP to nuclear bodies (Supplementary Figure S1 A-B), complementing our previous biochemical data^17^. Additionally, we observed that the fusion proteins sequestered endogenous NPC components: NUP88 (Figure 2D) and NUP62 (Figure 2E), both of which interact with NUP214 in NPCs, are recruited to SET-NUP214 nuclear foci in LOUCY cells. Similarly, NUP88 and NUP62 co-localized with SET-NUP214-GFP foci in transfected HCT-116 cells (Supplementary Figure S1 C-D). Interestingly, we also found that NUP98, the main component of the NPC barrier, co-localizes with the nuclear foci of SET-NUP214 (Figure 2E, Supplementary Figure S1E). Similar, albeit to a lesser extent, co-localization of NUP88 (Figure 2D), NUP62 (Figure 2E), and NUP98 (Figure 2F) was observed for DEK-NUP214 in FKH-1 cells as well as in transfected HCT-116 cells (Supplementary Figure S1 C-D). Altogether, the results suggest that SET-NUP214 and DEK-NUP214 perturb nucleocytoplasmic transport by accumulating CRM1-RanGTP nuclear export complexes and by disturbing the localization of endogenous NPC components.

**Figure 2:**
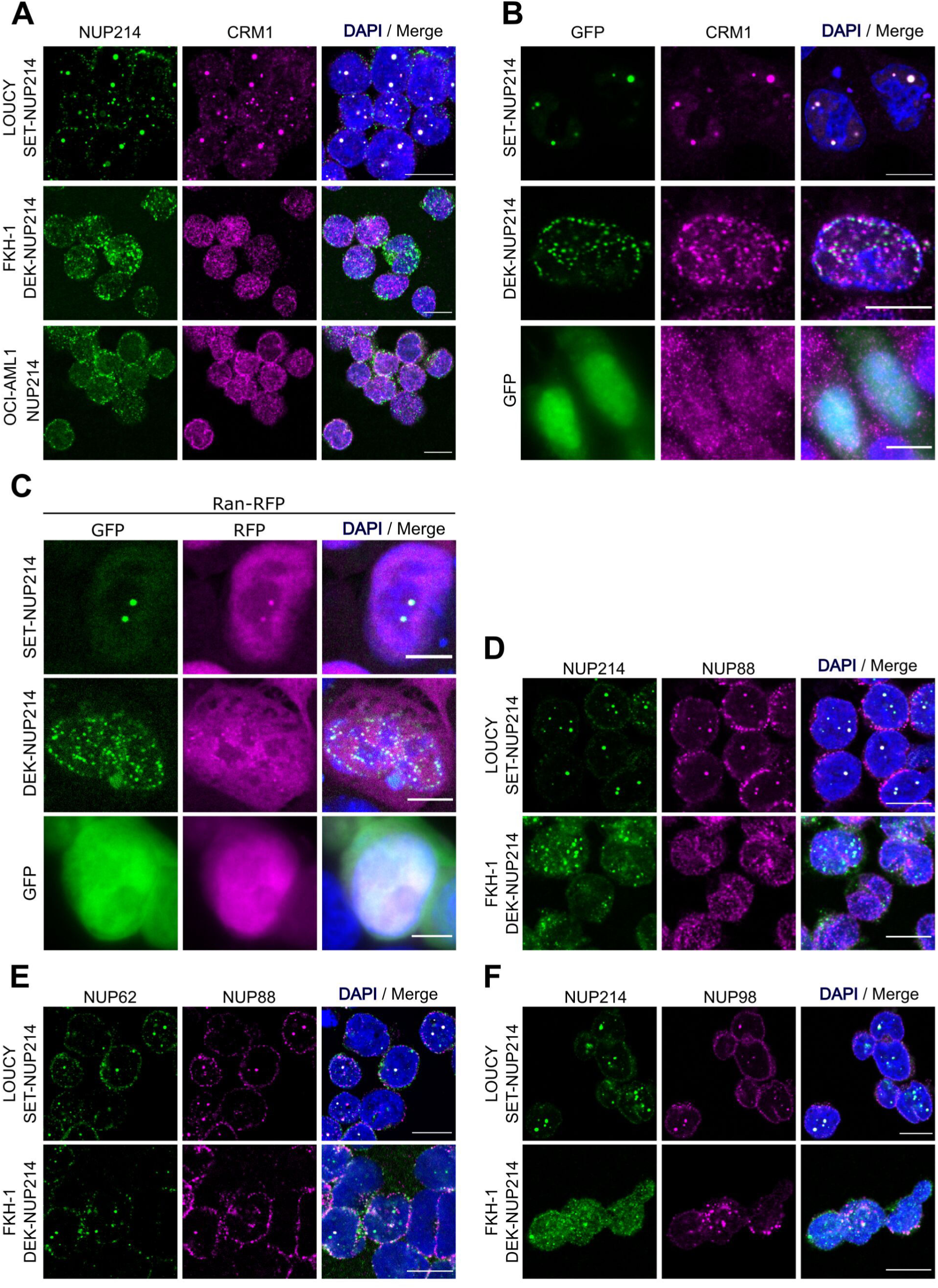
NUP214 fusions sequester the nuclear export factor CRM1 and endogenous nucleoporins to nuclei. **(A)** Cellular distribution of NUP214 (green) and CRM1 (magenta) in leukemia cell lines expressing either SET-NUP214 (LOUCY) or DEK-NUP214 (FKH-1). OCI-AML1 cells were used as control. **(B)** Co-localization of endogenous CRM1 (magenta) with SET-NUP214-GFP and DEK-NUP214-GFP fusion proteins (green) in transiently transfected HCT-116 cells. **(C)** Co-localization of SET-NUP214-GFP and DEK-NUP214-GFP with Ran-RFP in transiently transfected HCT-116 cells. **(D-F)** Colocalization of endogenous NUP88 (D), NUP62 (E) and NUP98 (F) with SET-NUP214 in LOUCY and DEK-NUP214 in FKH-1 cells. DNA was visualized with DAPI (blue). Shown are representative epifluorescence images. Scale bars, 10 µm.

### NUP214 fusion proteins are sensitive to nuclear export inhibition

CRM1 inhibition by LMB leads to the dissolution of nuclear bodies containing NUP214 fusion proteins, in LOUCY and in transfected cells^17, 18^. We confirmed these previous results for SET-NUP214 in LOUCY cells here (Figure 3 A-B, Supplementary Figure S1): treatment with LMB (20 nM, 3 h) resulted in a marked dispersal of SET-NUP214 nuclear bodies in LOUCY cells and a localization of SET-NUP214 and CRM1 throughout the nucleus (Figure 3 A-B, Supplementary Figure S2A). Similar results were obtained after the application of the SINE compound KPT-185 (1 µM, 24 h; Figure 3 A-B, Supplementary Figure S2A). Treatment with either LMB or KPT-185 led also to a significant decrease in DEK-NUP214 nuclear bodies in FKH-1 cells (Figure 3 C-D, Supplementary Figure S2A). In transfected cells, the application of LMB and KPT-185 similarly caused the dissolution of SET-NUP214-GFP and DEK-NUP214-GFP nuclear bodies and their homogenous distribution throughout the nucleoplasm, supporting the idea of an interplay between CRM1-mediated nuclear export and leukemia-related NUP214 fusion proteins (Supplementary Figure S2A). Detailed protocol used for quantification of nuclear bodies is presented in Supplementary Figure S2B.

**Figure 3:**
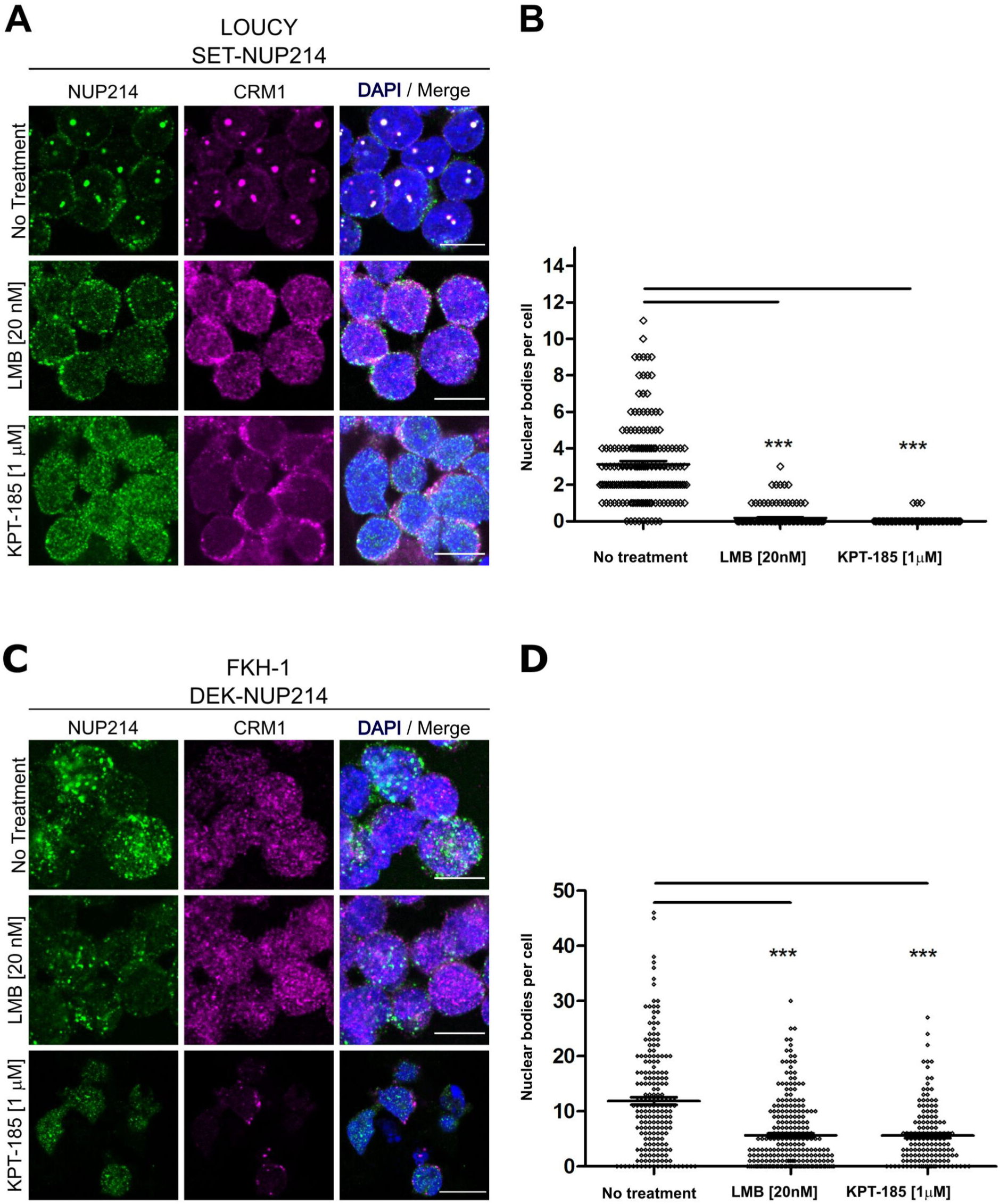
NUP214 fusion proteins are sensitive to inhibition of CRM1. LOUCY and FKH1 cells were treated with **(A)** LMB (20 nM, 3 h) or **(C)** KPT-185 (1 µM, 24 h) and NUP214 (green) and CRM1 (magenta) localization was studied by confocal microscopy. DNA stained with DAPI is depicted in blue. Scale bars, 10µm. Quantitative analysis of the number of NUP214 nuclear bodies in **(B)** LOUCY and **(D)** FKH-1 cells. At least 150 cells were analyzed for each condition. Statistical differences were calculated using GraphPad Prism (v5.01) using one-way analysis of variance (ANOVA) test. No treatment *vs* LMB; No treatment *vs* KPT-185. ***p<0.001

LMB binds irreversibly to CRM1, while KPT-185 interacts with CRM1 in a slowly reversible fashion^41^. We next examined the effect of drug withdrawal on nuclear body formation of SET-NUP214 in LOUCY cells. No prominent nuclear bodies were detectable immediately after the respective removal of LMB (Figure 4A, 0 h) and KPT-185 (Figure 4B, 0 h). Smaller SET-NUP214 nuclear bodies reformed 24 h after clearance from the drugs (Figures 4 A-B, 24 h), which were increasing to a similar size as those in non-treated cells within 48 h (Figures 4 A-B, 48 h and [-]). The re-appearance of SET-NUP214 nuclear bodies was accompanied by accumulation of CRM1 in these structures (Figure 5 A-B, 24 h and 48 h), supporting the hypothesis that SET-NUP214 localization and nuclear body formation depends on active CRM1.

**Figure 4:**
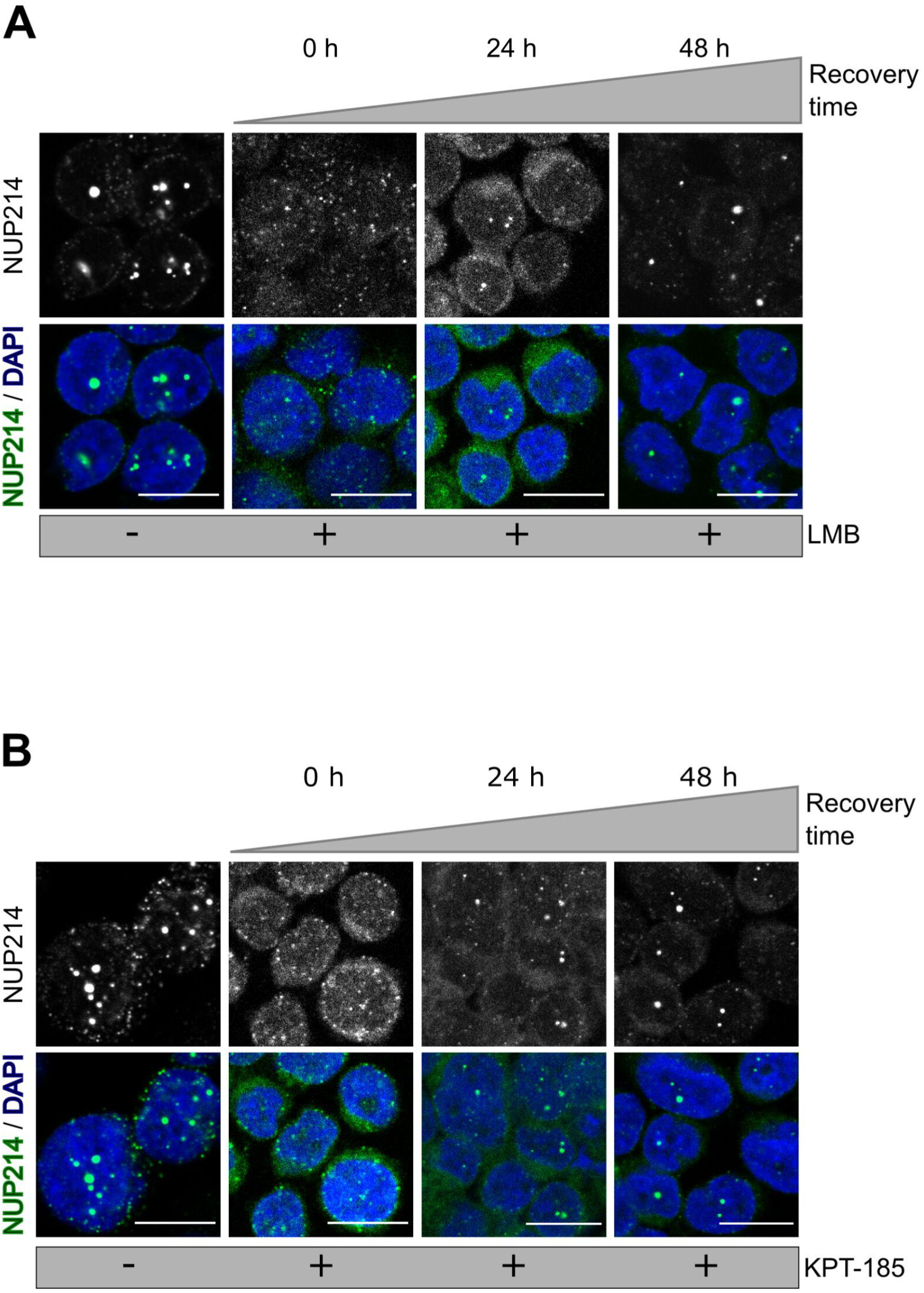
NUP214 nuclear foci reform after withdrawal of CRM1 inhibition. LOUCY cells (SET-NUP214) were treated with **(A)** LMB (20 nM, 3 h) or **(B)** KPT-185 (1 µM, 24 h). Cells were allowed to grow in drug-free medium for up to 48 h, after clearance from the respective drug. The presence of SET-NUP214 nuclear bodies was evaluated by immunofluorescence using anti-NUP214 (green) and anti-CRM1 (magenta) antibodies. DNA stained with DAPI is depicted in blue. Scale bars, 10µm

**Figure 5:**
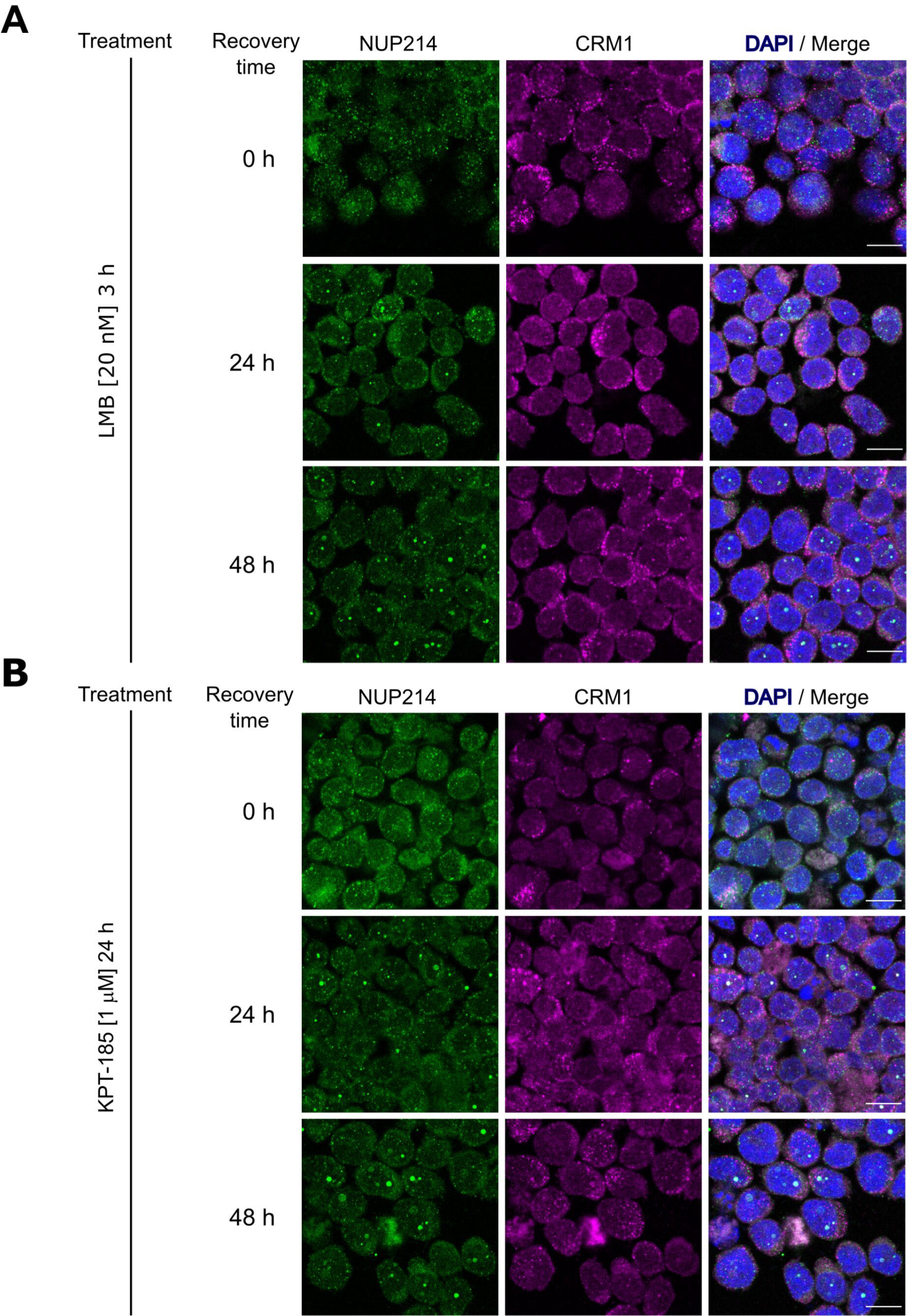
Formation of NUP214 nuclear after withdrawal of CRM1 inhibition is accompanied by CRM1 accumulation. LOUCY cells (SET-NUP214) were treated with **(A)** LMB (20 nM, 3 h) or **(B)** KPT-185 (1 µM, 24 h) and cells were allowed to grow in drug-free medium for up to 48 h after treatment. The formation of SET-NUP214 (green) nuclear foci is accompanied by the accumulation of CRM1 (magenta) in these structures. Shown are representative confocal images. DNA stained with DAPI is depicted in blue. Scale bars, 10 µm.

### Inhibition of CRM1-mediated nuclear export reduces cell viability

Next, we questioned whether CRM1 inhibition by LMB and KPT-185 affects the survival of leukemia cell lines harboring *NUP214* rearrangements. To address this question, LOUCY, MEGAL, and FKH-1 cells were treated with LMB or KPT-185 up to 72 h and cell viability was assessed at different time-points by Trypan blue exclusion dye assay. As shown in Figure 6 A-C, the viability of LMB- and KPT-185-treated LOUCY, MEGAL, and FKH-1 cells was significantly reduced as compared to untreated cells. Only about 47% of LOUCY (Figure 6A) and FKH-1 (Figure 6C) cells were alive 72 h after exposure to LMB, as compared to 77% of MEGAL cells (Figure 6B). The effect of KPT-185 on cell viability was slightly weaker in LOUCY (Figure 6A) and MEGAL (Figure 6B) cells as compared to LMB, but similar in FKH-1 cells (Figure 6C). Together these data suggest that CRM1 inhibition affects the viability of cells expressing NUP214 fusion proteins and that MEGAL cells appear to be more resistant to CRM1 inhibition in comparison to LOUCY and FKH-1 cells.

**Figure 6:**
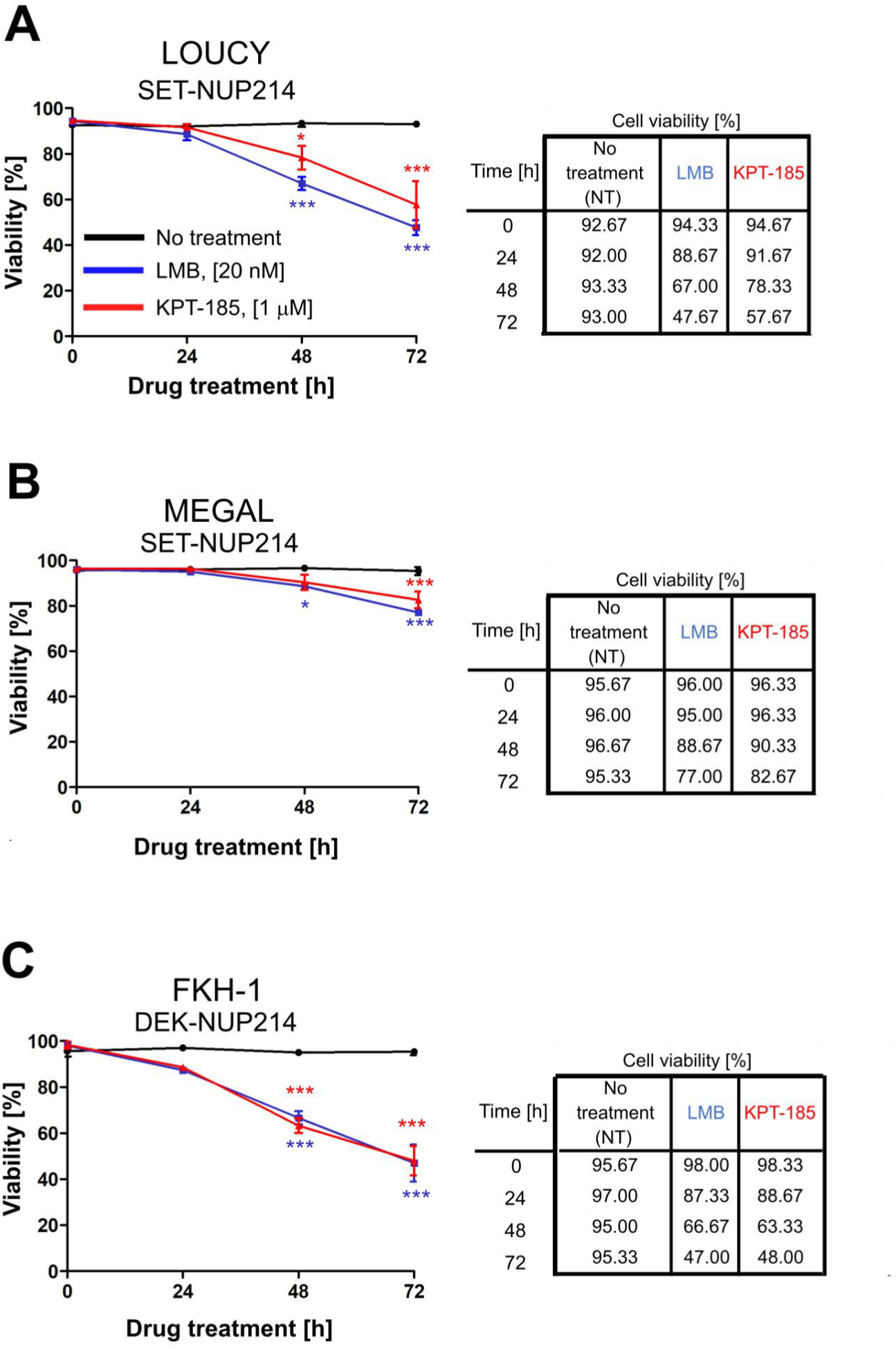
Inhibition of CRM1-mediated nuclear export reduces cell viability. Leukemia cell lines LOUCY (**A**), MEGAL (**B**), and FKH-1 (**C**) were treated with LMB (20 nM) or KPT-185 (1 µM) for the indicated time points and cell viability was measured by Trypan Blue exclusion dye. Statistical differences were calculated using GraphPad Prism (v5.01) using one-way analysis of variance (ANOVA) test. No treatment *vs* LMB; No treatment *vs* KPT-185. *p<0.05; **p<0.01; ***p<0.001.

### LMB and KPT-185 persistently affect cellular function

As mentioned above, LMB binds CRM1 irreversibly, in contrast to KPT-185^41^. This let us ask how cells respond to treatment withdrawal, i.e. how persistent are the effects of the two CRM1 inhibitors in NUP214-rearranged leukemia cells. We hence exposed the three leukemia cell lines to LMB or KPT-185 as described above and monitored cell viability and metabolic activity at different time points after drug removal. We found that after LMB and KPT-185 removal, cell viability (Figure 7A) and metabolic activity (Figure 7B) of MEGAL and FKH-1 cells were significantly reduced for at least 48 h. Similarly, LOUCY cells showed a significant reduction of cell viability for at least 48 h after drug removal, accompanied by a decrease in their metabolic activity, which nevertheless was not statistically significant. Again, the effect of the two CRM1 inhibitors on cell viability was weaker in MEGAL cell as in LOUCY and FKH-1 cells (Figure 7A; see also Figure 6). The persistent negative effect of CRM1 inhibition on cellular fitness of the leukemia cell lines was similarly observed in colony-forming assays (see Material and Methods) conducted with MEGAL and FKH-1 cells (Figure 7C). MEGAL cells formed colonies, which were smaller (< 50 µm) than colonies formed by non-treated cells (Figure 7C, top panel), whereas colonies formed by FKH-1 cells were reduced in size and number compared to non-treated cells (Figure 7C, bottom panel). These data further support the notion that MEGAL cells are slightly more resistant to CRM1 inhibition, when compared to LOUCY and FKH-1 cell lines. We additionally performed the experiment with LOUCY cells, however due to the small size of the colonies it was not possible to properly evaluate potential differences between treated and non-treated cells.

**Figure 7:**
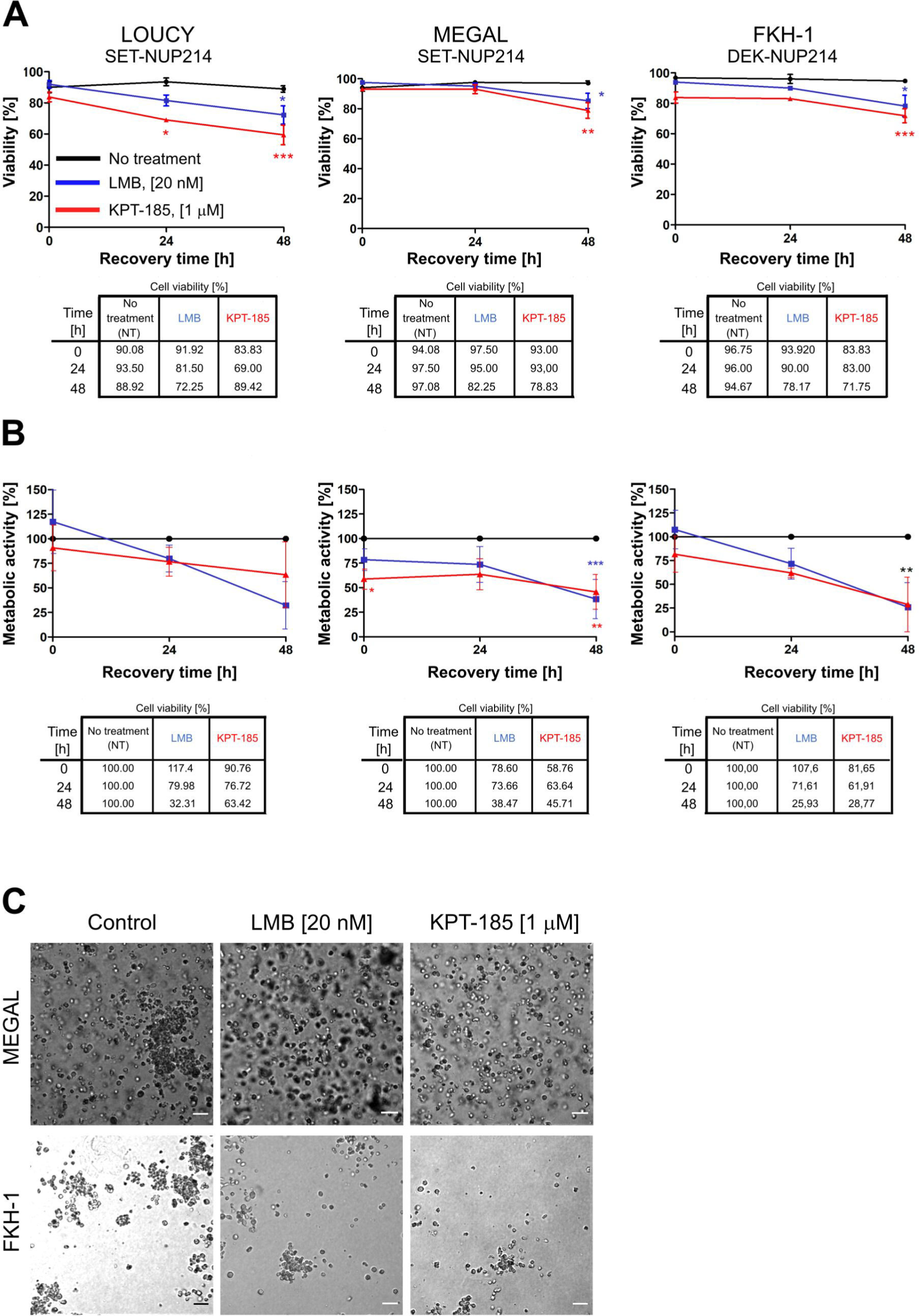
LMB and KPT-185 persistently affect cellular function. LMB or KPT-185 treated LOUCY, MEGAL, and FKH-1 cells were allowed to grow in drug-free medium for up to 48 h after drug clearance. **(A)** Cell viability was determined by Trypan Blue exclusion dye. **(B)** WST-1 assay was performed to measure cellular metabolic activity. Absorbance was measured at 450 nm. Results are normalized to untreated cells and expressed as percentage. **(C)** Colony forming assays of LMB and KPT-185 treated MEGAL and FKH-1 cells. Colonies were visualized under a 10x microscope objective after growth in drug-free Methocult™ medium for 14 days. Scale bars, 100 µm. Statistical differences were calculated using GraphPad Prism (v5.01) using one-way analysis of variance (ANOVA) test. No treatment *vs* LMB; No treatment *vs* KPT-185. *p<0.05; **p<0.01; ***p<0.001.

### CRM1 inhibition has varying effects on cell proliferation

We next examined the potential effect of CRM1 inhibitors on cell proliferation: enhanced proliferation is a hallmark of cancer cells. For that, we performed flow cytometry analysis of LOUCY, MEGAL, and FKH-1 cells exposed to LMB or KPT-185 and determined the number of proliferating cells based on the proliferation marker Ki-67 (Ki-67^+^ cells). In LOUCY cells, CRM1 inhibition by LMB had no significant impact on cell proliferation, with a maximum reduction of ∼11 %, 24 h after the treatment (Figure 8 A-B, left columns). Moreover, KPT-185 treatment had a significant (p<0.001) effect on cell proliferation, with a 27% reduction of Ki-67^+^ cells, 24 h after the treatment (Figure 8 A-B, left columns). Proliferation of MEGAL cells was not affected by LMB or KPT-185 treatment (Figure 8 A-B, middle columns), whereas FKH-1 cell proliferation was significantly lower upon LMB (p<0.05) and KPT-185 (p<0.01) treatment. Proliferation of LOUCY and FKH-1 cells was restored 48 h after KPT-185 removal. Proliferation of LMB-treated FKH-1 cells further decreased 48 h after drug removal, but also non-treated cells were less proliferative.

**Figure 8.**
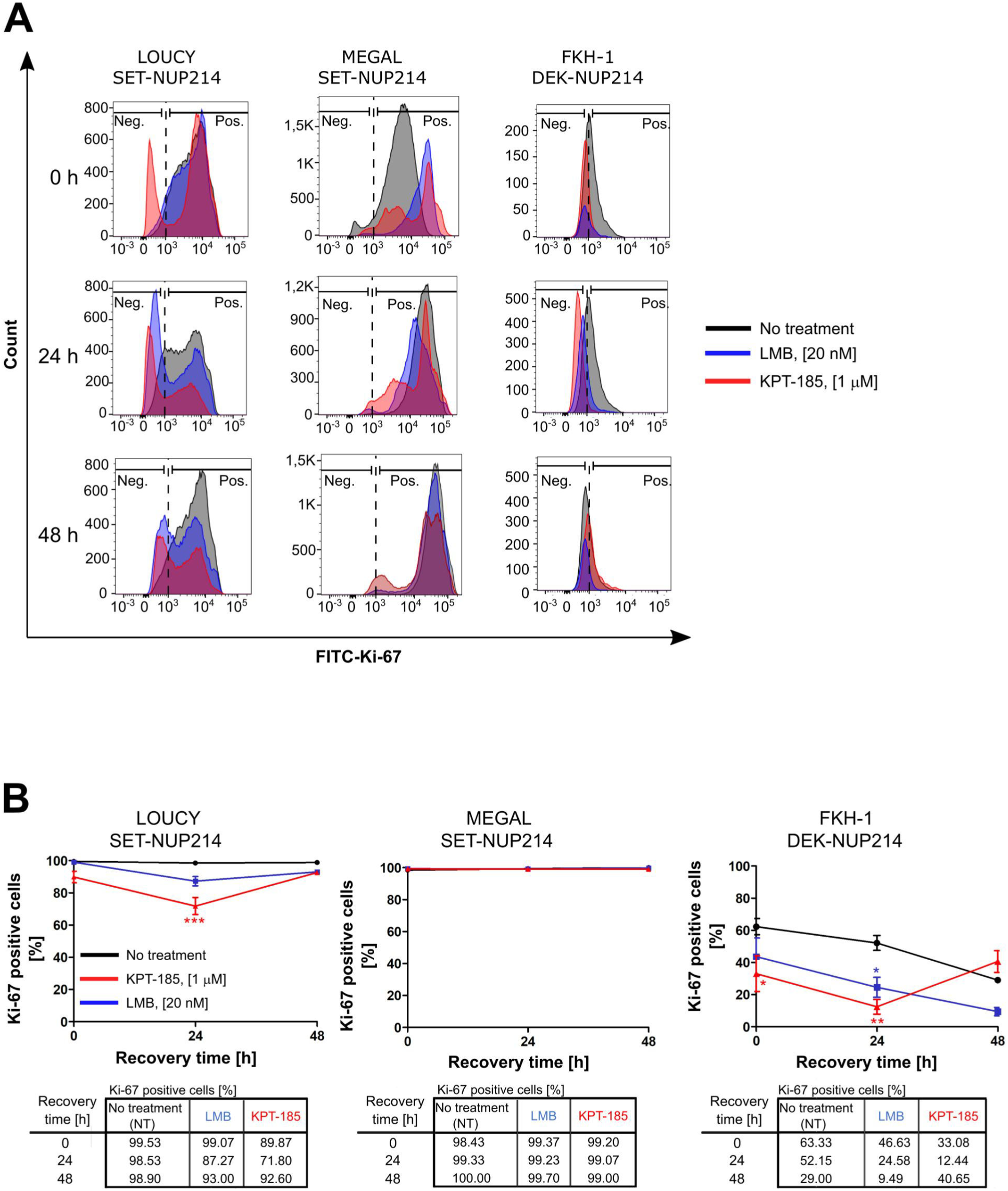
Distinct effects of CRM1 inhibition results on cell proliferation. Flowcytometric analysis of LOUCY, MEGAL, and FKH-1 cells after withdrawal from LMB or KPT-185 treatment. Cell proliferation was evaluated by using FITC-Ki-67 antibodies. **(A)** Histogram representation and **(B)** quantification of FITC-Ki-67 positive cells in the population of single cells at the indicated time points.

## Discussion

The pathologic potential of NUP214 fusion proteins has been widely recognized, but the underlying molecular mechanisms have been only sparsely studied^17, 19, 42-44^. NUP214 is an important player in nuclear export mediated by the ß-karyopherin CRM1 and CRM1 inhibition in turn has been proven beneficial in clinical trials as anti-cancer strategy. Here we confirm that endogenous and exogenous SET-NUP214 and DEK-NUP214 form nuclear bodies that accumulate CRM1 and its co-factor RanGTP (Figure 1 A, C, 2A-C, Supplementary Figure S1 A-B). Moreover, both NUP214 chimeras interfere with the cellular distribution of endogenous nucleoporins (Figure 2 D-F, Supplementary Figure S1 C-E), which might have a broad effect on nucleocytoplasmic trafficking. Further in-depth studies would be required to address this possibility.

In agreement with the idea of an interplay between NUP214 fusion proteins and CRM1-mediated nuclear export, two well-known CRM1 inhibitors, leptomycin B (LMB) and the selective nuclear export inhibitor (SINE) KPT-185, led to the dissolution of the SET-NUP214 and DEK-NUP214 nuclear bodies and their homogenous distribution throughout the nucleoplasm (Figure 3 A-D, Supplementary Figure S2). It remains to be established whether NUP214-fusion proteins sequester and trap CRM1 nuclear export complexes, or if CRM1 functions as a scaffold for the assembly of the fusion proteins into nuclear bodies. In any case, the disruption of these structures upon CRM1 inhibition indicates that NUP214 fusion proteins require CRM1 activity for nuclear body formation and possibly for their oncogenic capacities. Consistently, treatment of NUP214-rearranged leukemia cell lines with LMB or KPT-185 affects viability, metabolism, and proliferation, albeit to a different extent (Figure 6-8). This effect is sustained even after drug removal (Figure 7), likely to the stability in drug binding to CRM1. Binding of LMB and KPT-185 to CRM1 is energetically favored, i.e. both drugs not only bind free CRM1, but they also disrupt existing interactions between CRM1 and its cargoes^41^. Binding of LMB to CRM1 results in a covalent, irreversible interaction, whereas multiple hydrophobic bridges between KPT-185 and CRM1 result in a slowly reversible interaction^41^.

MEGAL cells respond different to CRM1 inhibition compared to LOUCY (and FKH-1) cells, i.e. they appear to be more robust/resistant (Figure 6-8). These discrepancies between MEGAL and LOUCY cells, which both express SET-NUP214, may be explained by differences in CRM1 levels in these two cell lines. Indeed, preliminary data indicate that in particular KPT-185 treatment coincided with reduced CRM1 levels in LOUCY cells and to a lesser extent in MEGAL cells and was also sustained after drug removal although some recovery could be observed (Supplementary Figure S3 A-B). LMB treatment on the contrary appears to not alter CRM1 protein levels. Further detailed analysis are required here, but our results are in accordance with previous reports which have shown that KPT-185 treatment led to reduced CRM1 protein levels, but not so LMB^20, 45-48^.

Cellular response to CRM1 inhibitors in leukemia has moreover been reported to depend on the mutational status of the tumor suppressor *TP53*^21^. LOUCY cells harbor a missense mutation p53^V272M^ (Supplementary Figure S4A), which results in functionally inactive p53^49, 50^. MEGAL cells feature a heterozygous frameshift mutation in *TP53* due to deletion of cytosine at position 898, resulting in a frameshift after a leucine residue at position 299, which affects p53’s 42 C-terminal residues (c.del898C, p53^299fs*42^; Supplementary Figure S4A). This mutant form of p53 in MEGAL cells is comparatively more abundant than the wild-type protein (Supplementary Figure S4B). The biological implications of p53^299fs*42^ remain to be elucidated. However, one may hypothesize that p53^299fs*42^ acts as dominant negative and hinders the biological function of the wild type counterpart, which may be stabilized by CRM1 inhibition^51, 52^. Such a dominant-negative effect of p53^299fs*42^ may contribute to the reduced sensitivity of MEGAL cells to CRM1 inhibition. Such a reduced sensitivity may arise from a reduced nuclear accumulation of tumor suppressor genes, delayed cell cycle, and/or reduced variation in overall protein expression after CRM1 inhibition, as seen in a fibrosarcoma cell line treated with the SINE KPT-330^53^. However, future studies are required to further elucidate this possibility.

Taken together, CRM1 inhibition might be an interesting therapeutic option in NUP214-related leukemia, similarly as described for several cancer models, including other forms of leukemia^23, 24, 29, 54, 55^. This notion is further supported by a recent study that showed successful disease remission for an AML patient with NUP214 driven leukemia who was treated with KPT-330 as single agent^56, 57^. This is particularly important, since no specific or targeted therapy has been developed so far^56^.

## Acknowledgements

We thank Drs. Denis Lafontaine and Martin Ruthardt for sharing reagents.

Confocal images were acquired at the CMMI, which is supported by the European Regional Development Fund (ERDF).

This work was supported by grants from the Fonds National de la Recherche Scientifique (F.R.S.–FNRS–FRIA, FC 16752) and Fondation Rose et Jean Hoguet to AM, as well as by research grants from the FNRS (T.0082.14 and J.013616F) and the Fédération Wallonie-Bruxelles (ARC 4.110. F.000092F) to BF.

