## Supplemental Information for "Targeted CRM1-inhibition perturbs leukemogenic NUP214 fusion proteins and exerts anti-cancer effects in leukemia cell lines with *NUP214* rearrangements"

#### Online Supplementary Methods

#### Cell lines and culture conditions

LOUCY (T-cell leukemia, ACC 394), MEGAL (acute megakaryoblastic leukemia, ACC 719), FKH-1 (acute myelocytic leukemia, ACC 614), OCI-AML1 (acute myeloid leukemia, ACC 726), and MOLM-13 (acute myeloid leukemia, ACC 554) were obtained from the Leibniz Institute, German Collection of Microorganisms and Cell Cultures GmbH (DSMZ; Braunschweig, Germany). HCT-116 cells were a gift from the Dr. Denis Lafontaine (Institute of Molecular Biology and Medicine, Université Libre de Bruxelles, Gosselies, Belgium). LOUCY, MEGAL, FKH-1, and MOLM-13 were cultured in RPMI-1640 medium (Gibco/Invitrogen; Merelbeke, Belgium) supplemented with 20% fetal bovine serum (FBS) and 1% penicillin/streptomycin (P/S; Gibco/Invitrogen). OCI-AML1 cells were cultured in α-MEM medium (LONZA$^{TM}$ BioWhittaker$^{TM}$; Verviers, Belgium) supplemented with 20% FBS and 1% P/S. For FKH-1 and OCI-AML1 cells, the medium was further supplemented with 0.1 ng/ml recombinant human growth-colony stimulating factor (G-CSF; PREPROTECH, London, UK). HCT-116 cells were cultured in McCoy's 5A medium (LONZA$^{TM}$ BioWhittaker$^{TM}$), supplemented with 10% FBS and 1% P/S. All cell lines were kept in a humidified incubator at 37ºC with 5% $CO_2$ atmosphere.

#### Generation of plasmids

##### SET-NUP214

For the cloning of SET-NUP214-GFP, total RNA ($RNA_T$) was extracted from LOUCY cells, which carry del(6)(q23) resulting in the *SET-NUP214* fusion transcript, using the High Pure RNA Isolation Kit (Roche Life Sciences, Basel, Switzerland) according to the manufacturer's instructions and stored at -80°C for further use. The synthesis of cDNA was performed by reverse transcription-polymerase chain reaction (RT-PCR) as follows: 100 ng of $RNA_T$ were mixed with 3,4 μg of deoxy-thymidine oligomer $(OligodT)_{12-18}$ primer and 10 nmol of desoxyribonucleotides (dNTPs) to a final volume of 20 μl and incubated at 65°C for 5 min. The RT-PCR reaction was initiated by the addition of 2 μl of 0,1 μM DTT, 4 μl of 5x First Strand Buffer, 40 units (U) of RNase OUT™ and 200 U of SuperScript™ Reverse Transcriptase II (SSII-ThermoFisher Scientific, Merelbek, Belgium). The final mixture was incubated at the following thermal program: 42°C for 2 min, 25°C for 10 min and 42°C for 1 h. The resulting cDNA was immediately used for PCR amplification or stored at -20°C for further applications. Next, the converted cDNA was used as template for the amplification of the *SET-NUP214* coding sequence by PCR using specific primers (forward 5'- ATGTCGGCGCCGGCGGCCAA-AGTCAGTAAA-3'; reverse 5'- TATCCCGGGCTTCGCCAGCCACCA-AAACC-3'). The resulting PCR product was purified using the High Pure PCR Product Purification Kit (Roche Life Sciences) according to manufacturer's instructions and inserted into the pGEM®-T intermediate vector backbone (Promega, Leiden, Netherlands). The ligation product was used to transform XL1-Blue competent cells and positive colonies were selected by blue/white screening. Plasmid DNA was isolated using the GenElute™ Plasmid MidiPrep Kit (Sigma Aldrich, Overijse, Belgium), according to manufacturer's instructions. To generate the SET-NUP214-GFP construct, the SET-NUP214 coding sequence was subcloned from the pGEM-T backbone into the pEGFP-N3 vector, using the SacI and SalI restriction enzymes (New England Biolabs – NEB; Ipswich, Massachusetts, USA). Digestion products were purified using the High Pure PCR Product Purification Kit (Roche Life Sciences) according to manufacturer's instructions. Ligation was performed for 2 h using T4 DNA ligase (Thermo Fisher Scientific) and the purified ligation product was transformed into XL1-blue competent cells. Cells were then seeded on LB-agar plates supplemented with kanamycin (50 µg/ml) and grown overnight at 37ºC. Colonies were screened by colony-PCR and positive clones were expanded in LB/kanamycin medium. The pEGFP-SET-NUP214 plasmid was fully sequenced to verify the correct sequence.

**DEK-NUP214-GFP**

The pENTR1-DEK-NUP214 Gateway entry vector was a gift from Dr Martin Ruthardt (Cardiff University, UK). The DEK-NUP214 coding sequence was subcloned into the peZY-EGFP destination vector using the Gateway™ LR clonase™ enzyme mix (Invitrogen), according to the manufacturer's instructions. The peZY-EGFP destination vector encodes the toxic *ccdb* gene. Upon the LR clonase reaction, the *ccdb* gene is replaced by the DEK-NUP214 sequence, rendering the plasmid non-toxic. For transformation, the *ccdb*-sensitive TOP10 competent cells were used. The peZY-EGFP-DEK-NUP214 plasmid was fully sequenced to verify the correct sequence.

pcDNA-RanWT-mRFP1-polyA, pcDNA-Ran-Q69L-mRFP1-polyA and pcDNA-Ran-T24N-mRFP1-polyA were a gift from Yi Zhang (Addgene plasmid #59750, #104560 and #104561, respectively; (Inoue et al. 2014)).

All transfections were carried out for 48 h and performed using the JetPrime Transfection reagent (Polyplus Transfection; city, country), according to the manufacturer's instructions.

**Immunofluorescence of suspension cells**

Leukemia cell lines were seeded at $0.8x10^6$ cells/ml and grown for 24 h - 48 h. Cells were fixed at a concentration of $1×10^6$ cells/ml in 2% formaldehyde for 15 min. Next, cells were washed twice for 10 min in PBS and permeabilized with PBS containing 2% bovine serum albumin (BSA) and 0.1% Triton X-100 for 20 min. Cells were then resuspended in 50 µl of the appropriate primary antibodies diluted in PBS containing 2% BSA and incubated overnight at 4°C. Cells were washed twice with PBS/2% BSA/0.1% Triton X-100, resuspended in 50 µl of the appropriate secondary antibodies diluted in PBS/2% BSA, incubated for 1 h at room temperature and washed again twice in PBS/2% BSA/0.1% Triton X-100. Next, cells were resuspended in 50-100 µl of PBS, seeded on polylysine-coated glass slides, and incubated at 37ºC until dry. Slides were mounted with Mowiol-4088 (Sigma-Aldrich) containing DAPI (1 µg/ml) and stored at 4°C until viewed.

**Immunofluorescence of adherent cells**

HCT-116 were grown on polylysine-coated glass coverslips and fixed in 2% formaldehyde for 15 min, washed three times for 10 min with PBS, and permeabilized with PBS/2% BSA /0.1% Triton X-100 for 10 min. Next the cells were washed twice for 10 min in PBS/2% BSA, incubated with the appropriate primary antibodies overnight, washed twice in PBS/2% BSA/0.1% Triton X-100, incubated with the appropriate secondary antibodies for 1 h, washed twice for 10 min with PBS, mounted with Mowiol-4088 containing DAPI (1 µg/ml), and stored at 4°C until viewed.

Cells were imaged using a Zeiss LSM-710 (Zeiss, Oberkochen, Germany) confocal laser-scanning microscope. Images were recorded using the microscope system software and processed using ImageJ (http://imagej.nih.gov) and Inkscape 0.92 Software (https://inkscape.org).

#### Antibodies

The following antibodies were used for immunofluorescence microscopy: rabbit polyclonal anti-NUP214 (abcam, Cambridge, UK, ab740497; dilution 1:500), mouse monoclonal anti-XPO1 (BD Transduction, San Jose, CA 611832; dilution 1:200), mouse monoclonal anti-NUP88 (BD Transduction, dilution 1:500), rabbit polyclonal anti-NUP62 (Santa Cruz Biotechnology, Heidelberg, Germany, dilution 1:100) and rat anti-NUP98 (Sigma-Aldrich, N1038, dilution 1:1000). Alexa Fluor-488 and Alexa Fluor-568 (Invitrogen/ThermoFisher, dilution 1:1000) conjugates were used as secondary antibodies.

The following antibodies were used for Western blotting: rabbit polyclonal anti-NUP214 (abcam, ab740497, dilution 1:5000), mouse-monoclonal anti-p53 (Santa Cruz Biotechnology, dilution 1:500), mouse monoclonal anti-CRM1 (BD Transduction 611832; dilution: 1:500), rabbit polyclonal anti-actin (Sigma Aldrich, a2066, dilution 1:1000), rabbit polyclonal anti-α-tubulin (abcam, ab18250, dilution 1:4000). Alkaline phosphatase coupled antibodies (Sigma Aldrich, anti-rabbit IgG, a3687; anti-mouse IgG, a9316, dilution 1:10.000) were used as secondary antibodies.

#### Cell viability and proliferation count

Cells were seeded at $0.8 \times 10^6$ cells/ml 24 h before treatment with 20 nM LMB or 1 µM KPT-185 for 24 h, 48 h and 72 h. At each time point, cell number and viability were assessed by Trypan Blue exclusion dye assay. For recovery time experiments, cells were treated for 3 h with 20 nM LMB or for 24 h with 1 µM KPT-185. Cells were then washed and cultured in fresh culture medium for 0 h, 24 h or 48 h. At each indicated time point cell viability and cell number were measured by Trypan Blue exclusion dye assay.

**WST-1 assays**

Cells were seeded at $0.8 \times 10^6$ cells/ml 24 h before treatment with 20 nM LMB for 3 h or 1 µM KPT-185 h for 24 h. After the respective treatment, cells were washed and cultured in fresh culture medium for 0 h, 24 h or 48 h. WST-1 assays were performed using the Cell Proliferation Reagent WST-1 (Roche Life Sciences) according to the manufacturer's instructions. Absorbance (450 nm and 650 nm) was measured in triplicate in a SYNERGY™ Mx microplate reader (BioTek, Bad Friedrichshall, Germany) and the data were acquired with the GEN5 Data Analysis software (BioTek).

**Colony Forming Cell (CFC) assays**

Cells were seeded at $0.8 \times 10^6$ cells/ml 24 h before treatment with 20 nM LMB for 3 h or 1 µM KPT-185 for 24 h. After treatment, cells were washed and resuspended in 3 ml of MethoCult™ H4100 semisolid culture medium (STEMCELL Technologies, Cambridge UK), supplemented with 20% FBS and 1% P/S. For FKH-1 cells, the culture medium was additionally supplemented with 0.1 ng/ml G-CSF. Cells were maintained for 14 d and visualized at 100× amplification using a Zeiss Observer.Z1 (Zeiss) microscope. Images were recorded using the microscope system software and processed using ImageJ (http://imagej.nih.gov) and Inkscape 0.92 Software (https://inkscape.org).

**Ki-67 proliferation assays**

Cells were seeded at $0.8 \times 10^6$ cells/ml 24 h before treatment with 20 nM LMB for 3 h or 1 µM KPT-185 h for 24 h. Next, cells were washed and cultured in fresh culture medium for 0 h,

24 h or 48 h. At each timepoint, cells were harvested and fixed in 5 ml of ice-cold 70% ethanol. Cells were kept at -20ºC in 70% ethanol until further use. For Ki-67 staining, cells were washed twice in wash buffer (PBS complemented with 1% FBS, 0.09% sodium azide (NaN3), pH7.2), resuspended in wash buffer at $1\times10^7$ cells/ml and incubated with FITC-Ki-67 (BD Biosciences) primary antibody for 35 min in the dark, with agitation at room temperature. Next, cells were washed twice in wash buffer and resuspended in 500 µl of PBS for flow cytometry analysis. Flow cytometric acquisition was performed on a FACS Canto II machine (BD Biosciences). Data were processed using FlowJo V10 (Tree Star Inc.).

**Western Blot**

Cells were lysed in lysis buffer (50 mM Tris-HCl, pH 7.8, 150 mM NaCl, 1% Nonidet-P40 and protease inhibitor cocktail tablets (Roche, Basel Switzerland)). Bradford assay was used to determine protein concentration and 35 µg of protein were loaded and separated by sodium dodecyl sulfate-polyacrylamide gel electrophoresis (SDS-PAGE; 5% or 7%). The proteins were transferred onto a PVDF membrane (Immobilon-P, Millipore) and the membranes were blocked with TBS containing 0.1% Tween 20 and 5% non-fat dry milk for 1 h. The membranes were then incubated overnight in blocking solution containing a primary antibody followed by washing 3 times in TBS containing 0.1% Tween 20 and 5% non-fat dry milk. The membranes were next incubated with secondary alkaline phosphatase coupled antibodies for 1 h, washed 3 times in TBS and developed. X-ray films were scanned and processed using ImageJ.

***TP53* Sanger-sequencing**

Total RNA ($RNA_T$) was extracted from LOUCY and MEGAL cells, using the High Pure RNA Isolation Kit (Roche Life Sciences, Basel, Switzerland) according to the manufacturer's instructions and stored at -80°C for further use. The synthesis of cDNA was performed by RT-PCR as previously described, as described for the generation of the SET-NUP214 construct. Next, the converted cDNA from LOUCY cells was used as template for the amplification of the exon 7 of *TP53* by PCR using specific primers (forward 5'-CTACATGTGTAACAGTTCCTG-3' and reverse 5'-GAAATATTCTCCATCCAGTGG-3'). The cDNA from MEGAL cells was used as template for the amplification of the complete coding sequence of *TP53* using the specific primers: forward 5'- ATGGAGGAGCCGCAGTCA-3' and reverse 5'-GTATCAGGCAAAGTCATAGAACCA-3'. PCR reactions were performed using the Q5® High Fidelity DNA Polymerase (NEB) according to the manufacturer's instructions.

### Figure S1

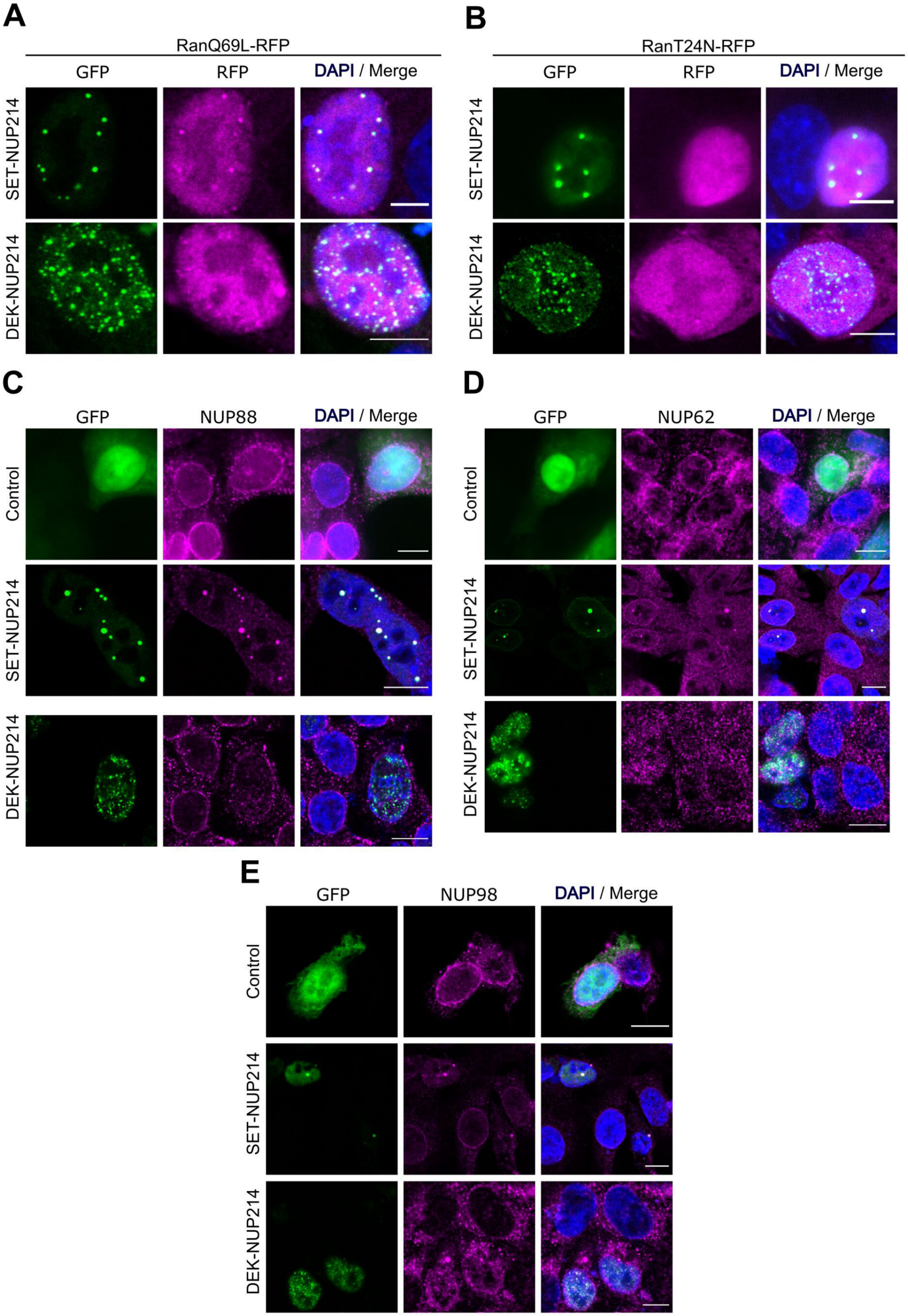
A
RanQ69L-RFP
GFP
RFP
DAPI / Merge
SET-NUP214
DEK-NUP214
B
RanT24N-RFP
GFP
RFP
DAPI / Merge
SET-NUP214
DEK-NUP214
C
GFP
NUP88
DAPI / Merge
Control
SET-NUP214
DEK-NUP214
D
GFP
NUP62
DAPI / Merge
Control
SET-NUP214
DEK-NUP214
E
GFP
NUP98
DAPI / Merge
Control
SET-NUP214
DEK-NUP214

### Figure S2

## A

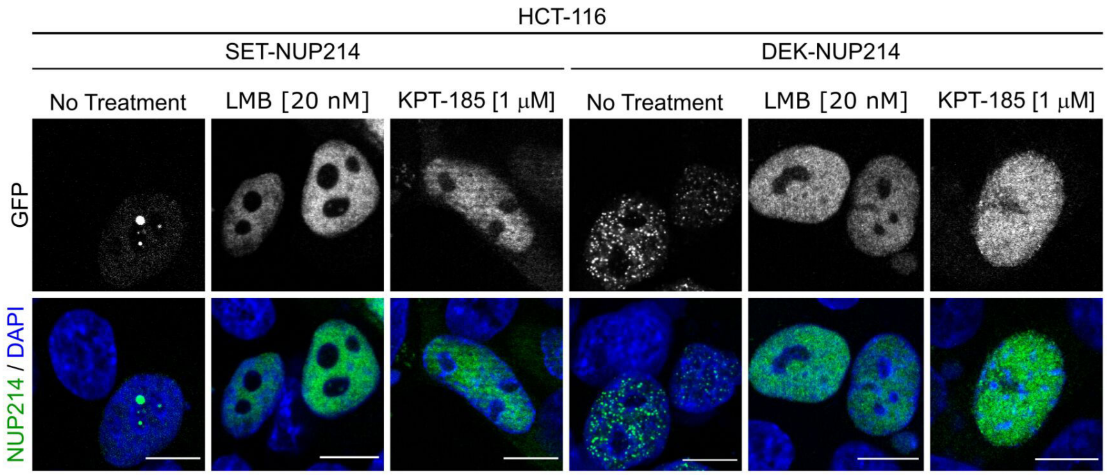

HCT-116
SET-NUP214
DEK-NUP214
No Treatment
LMB [20 nM]
KPT-185 [1 μM]
No Treatment
LMB [20 nM]
KPT-185 [1 μM]
GFP
NUP214 / DAPI


## B

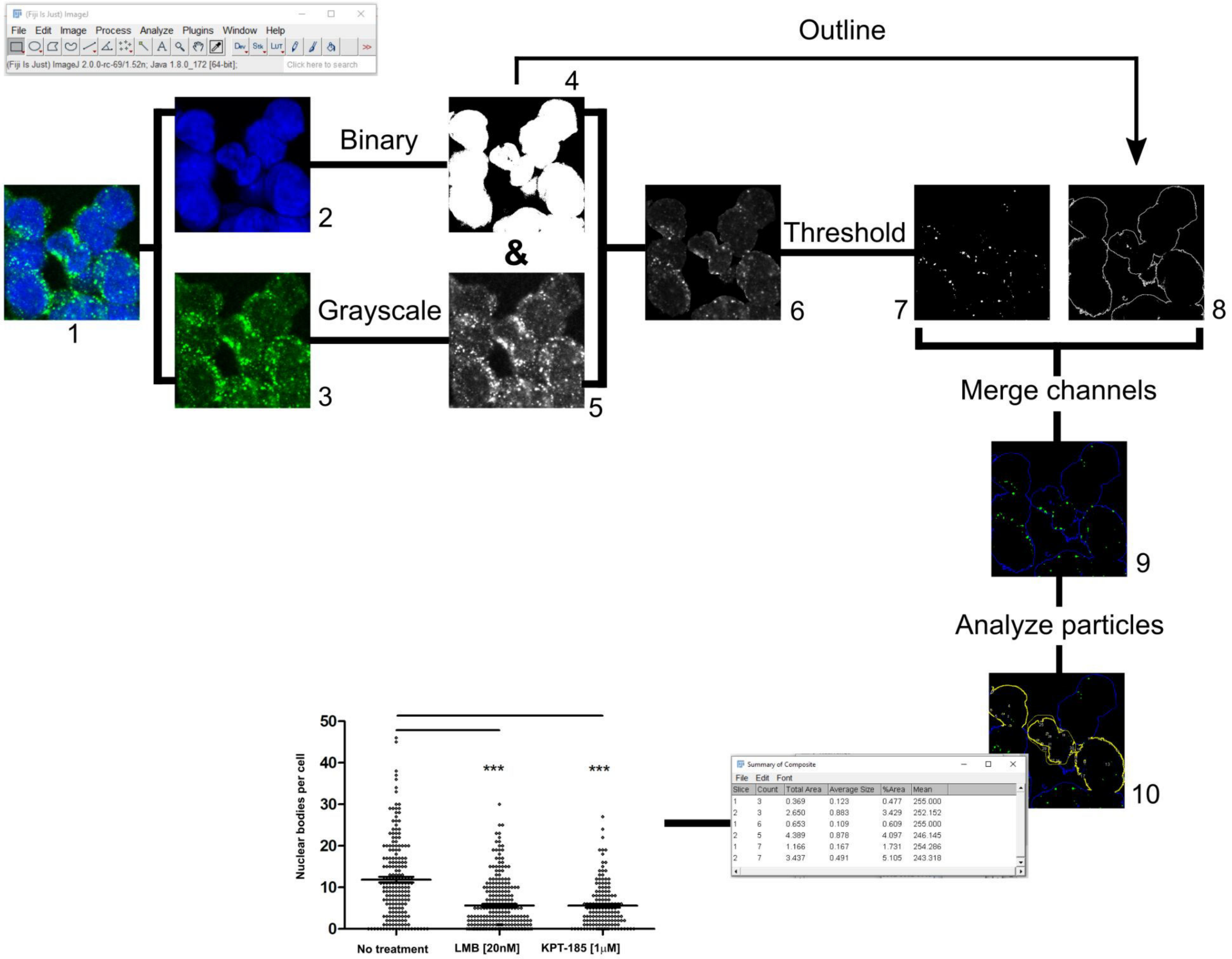

(Fiji Is Just) ImageJ
File Edit Image Process Analyze Plugins Window Help
(Fiji Is Just) ImageJ 2.0.0-rc-69/1.52n; Java 1.8.0_172 [64-bit];
Click here to search
Outline
Binary
&
Grayscale
Threshold
Merge channels
Analyze particles
1
2
3
4
5
6
7
8
9
10
Summary of Composite
File Edit Font
Slice Count Total Area Average Size %Area Mean
1 3 0.369 0.123 0.477 255.000
2 3 2.650 0.883 3.429 252.152
1 6 0.653 0.109 0.609 255.000
2 5 4.389 0.878 4.097 246.145
1 7 1.166 0.167 1.731 254.286
2 7 3.437 0.491 5.105 243.318
Nuclear bodies per cell
0
10
20
30
40
50
***
***
No treatment
LMB [20nM]
KPT-185 [1μM]

#### Figure S3

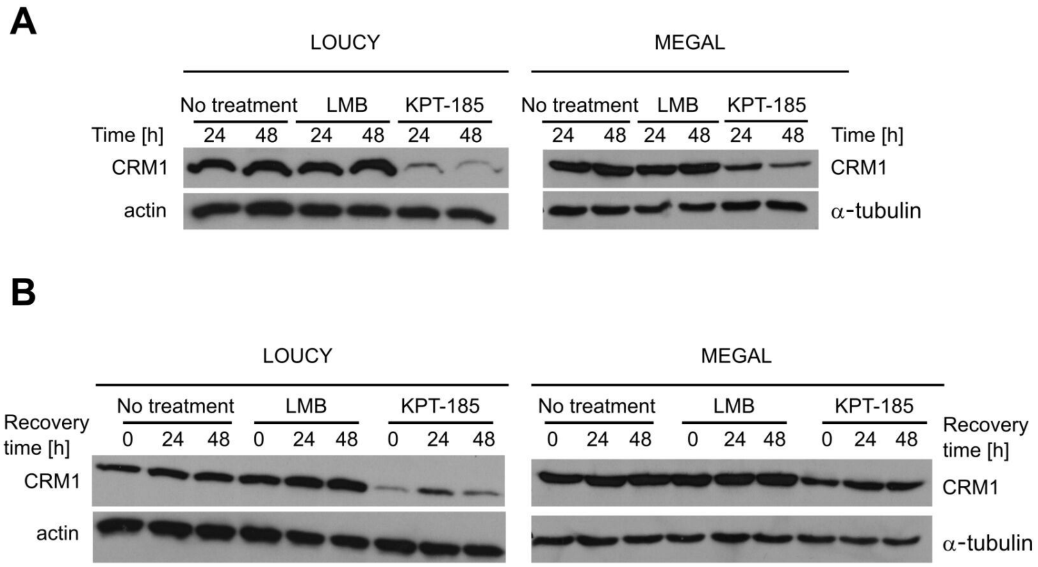
A
LOUCY
MEGAL
No treatment
LMB
KPT-185
Time [h]
24
48
CRM1
actin
α-tubulin
B
Recovery time [h]
0
24
48

**Figure S4**

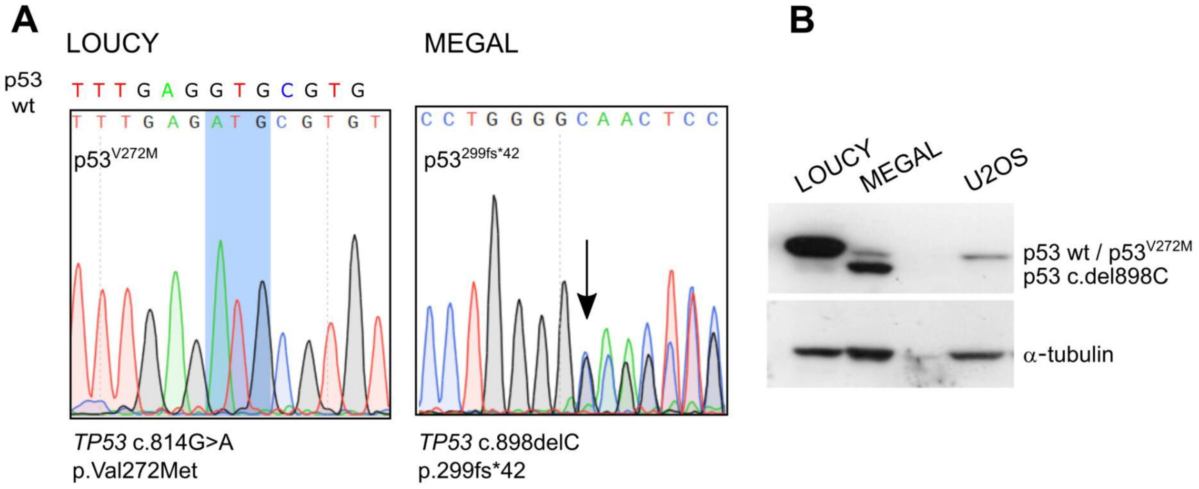
A
LOUCY
MEGAL
p53 wt
T T T G A G G T G C G T G
T T T G A G A T G C G T G T
p53V272M
C C T G G G G C A A C T C C
p53299fs*42
TP53 c.814G>A
p.Val272Met
TP53 c.898delC
p.299fs*42
B
LOUCY
MEGAL
U2OS
p53 wt / p53V272M
p53 c.del898C
α-tubulin

**Supplementary Figure S1:** NUP214 fusions accumulate the nuclear export factor CRM1 and perturb the localization of endogenous nucleoporins. NUP214 fusion proteins (green) accumulate **(A)** RFP-tagged RanQ69L (magenta) to nuclear foci, but not **(B)** RFP-tagged mutant RanT24N (magenta). Expression of SET-NUP214-GFP and DEK-NUP214-GFP (green) perturbs the localization of endogenous **(C)** NUP88, **(D)** NUP62, and **(E)** NUP98 and cause their accumulation to nuclear foci formed by NUP214 fusion proteins. Shown are representative confocal images. DNA stained with DAPI is depicted in blue. Scale bars 10 μm.

**Supplementary Figure S2:** NUP214 fusion proteins are sensitive to CRM1 inhibition. **(A)** HCT-116 cells were transfected with SET-NUP214-GFP or DEK-NUP214-GFP and treated with LMB or KPT-185. Nuclear foci formation was visualized GFP fluorescence, DNA by DAPI staining. Shown are representative confocal images. Scale bars 10 μm. **(B)** Schematic representation of the ImageJ protocol used to quantify nuclear foci formation in LOUCY and FKH-1 cells after treatment with LMB or KPT-185. Control and treated cells were analyzed by confocal microscopy after staining with an anti-NUP214 antibody. Composite images (1) were split into 358 nm (DAPI, nucleus, 2) and 488 nm (NUP214, 3) channels. Image 2 was converted to the corresponding binary (4), and image 3 was converted to grayscale (5). Then, image 5 was subtracted from image 4, which resulted in an image of the NUP214 fluorescence contained in the nuclear area (6). After, image 4 was converted to the corresponding outline (8), to delineate individual nuclei. A fluorescence threshold was applied to image 6 maintain only the signal corresponding to the nuclear foci (7). After, images 7 and 8 were merged and particles were counted in each individual nucleus. The results were recorded and exported to Excel files. Graphs were generated with the statistical analysis software GraphPad Prism.

**Supplementary Figure S3:** LOUCY and MEGAL cell lines respond differently to CRM1 inhibition by KPT-185. Variation of CRM1 protein levels in LOUCY and MEGAL cell lines after treatment with LMB and KPT-185. **(A)** Cells were treated with 20 nM LMB and 1 μM Kpt-185 for up to 48 h. **(B)**: Cells were treated with 20 nM LMB for 3 h and 1 μM Kpt-185 for 24 h. After, cells were washed and cultured in drug-free medium and allowed to recover for up to 48 h.

**Supplementary Figure S4:** *TP53* mutation analysis in LOUCY and MEGAL cell lines. **(A)** Sanger sequencing of *TP53* using cDNA from LOUCY and MEGAL cell lines. LOUCY cells harbor a homozygous or hemizygous missense mutation (c.814G>A) that results in the synthesis of a mutant p53 ($p53^{V272M}$). MEGAL cells harbor a deletion (c.898delC) that results in a frameshift, leading to the formation of a mutant form of p53 ($p53^{299fs*42}$). **(B)** p53 expression in LOUCY and MEGAL cell lines by western blot. A single band is detected in LOUCY cells, which corresponds to $p53^{wt}$, $p53^{V272M}$ or both. MEGAL cells express the two p53 variants, with higher expression of the mutant $p53^{299fs*42}$ form than $p53^{wt}$.
